# Functional connectivity-based subtypes of individuals with and without autism spectrum disorder

**DOI:** 10.1101/198093

**Authors:** Amanda K. Easson, Zainab Fatima, Anthony R. McIntosh

**Affiliations:** Rotman Research Institute, Baycrest Hospital, Toronto, ON, Canada; Department of Psychology, University of Toronto, Toronto, ON, Canada; Department of Psychology, Faculty of Health, Sherman Health Sciences Centre, York University, Toronto, ON, Canada

**Keywords:** autism spectrum disorder, functional connectivity, clustering, brain-behaviour relationships, multivariate statistics, resting-state networks

## Abstract

Autism spectrum disorder (ASD) is a heterogeneous neurodevelopmental disorder, characterized by impairments in social communication and restricted, repetitive behaviours. Neuroimaging studies have shown complex patterns of functional connectivity (FC) in ASD, with no clear consensus on brain-behaviour relationships or shared patterns of FC with typically developing controls. Here, we used k-means clustering and multivariate statistical analyses to characterize distinct FC patterns and FC-behaviour relationships in participants with and without ASD. Two FC subtypes were identified by the clustering analysis. One subtype was defined by increased FC within resting-state networks and decreased FC across networks compared to the other subtype. A separate FC pattern distinguished ASD from controls, particularly within default mode, cingulo-opercular, sensorimotor, and occipital networks. There was no significant interaction between subtypes and diagnostic groups. Finally, analysis of FC patterns with behavioural measures of IQ, social responsiveness and ASD severity showed unique brain-behaviour relations in each subtype, and a continuum of brain-behavior relations from ASD to controls within one subtype. These results demonstrate that distinct clusters of FC patterns exist in both ASD and controls, and that FC subtypes can reveal unique information about brain-behaviour relationships.

**Author Summary:** Autism spectrum disorder (ASD) is a neurodevelopmental disorder, with high variation in the types of severity of impairments in social communication and restricted, repetitive behaviours. Neuroimaging studies have shown complex patterns of communication between brain regions, or functional connectivity (FC), in ASD. Here, we defined two distinct FC patterns and relationships between FC and behaviour in participants with and without ASD. One subtype was defined by increased FC within distinct networks of brain regions, and decreased FC between networks compared to the other subtype. A separate FC pattern distinguished ASD from controls. The interaction between subtypes and diagnostic groups was not significant. Analysis of FC patterns with behavioural measures revealed unique information about brain-behaviour relations in each subtype.

## INTRODUCTION

Autism spectrum disorder (ASD) is a neurodevelopmental disorder that is characterized by impairments in social cognition as well as restricted and repetitive behaviours (RRBs; American Psychiatric Association, 2013). ASD is a highly heterogeneous disorder, with a broad range of the types and severities of behaviours that can be displayed. For instance, verbal and nonverbal IQ are highly variable in ASD (e.g. Munson et al., 2008), and RRBs can range from low-level motor stereotypies to higher-order behaviours such as insistence on sameness (American Psychiatric Association, 2013). It has been proposed that these complex behavioural features are associated with atypical patterns of functional connectivity (FC). Such theories include reduced communication between frontal and posterior brain regions (Just et al., 2012), increased local FC along with reduced long-range FC (Belmonte et al., 2004; Courchesne & Pierce, 2005), and an abnormal developmental trajectory of FC compared to typically developing (TD) individuals (Nomi & Uddin, 2015; Uddin et al., 2013b). However, complex patterns of both increased and decreased FC have been found in neuroimaging studies of ASD, and results are inconsistent across studies (see Hull et al., 2016; Picci et al., 2016; and Uddin et al., 2013b for reviews).

It is crucial to consider the heterogeneous nature of ASD, both in terms of behavioural severity and FC profiles. The importance of this consideration is highlighted by the inconsistent results regarding relationships between FC and behavioural profiles in individuals with ASD in previous studies (e.g. Keown et al., 2013; Lee et al., 2016; Monk et al., 2009; Uddin et al., 2013b). Several recent studies that considered the heterogeneity of neurobiological and behavioural features of ASD have reported novel finding regarding brain-behaviour relationships. For instance, Hahamy, Behrmann & Malach (2015) found that idiosyncratic distortions in FC from a “typical” template were related to ASD symptom severity. Nunes et al. (2018) also reported that incorporation of vertices along the cortical surface into intrinsic connectivity networks, particularly into default mode and sensorimotor networks, was more idiosyncratic in ASD and related to ASD symptom severity.

Defining subtypes of ASD based on FC metrics has the potential to resolve some of the current discrepancies in the literature regarding the nature of FC abnormalities in individuals with this disorder, as well as to shed light on the complex relationships between FC and behaviour, which may differ between subtypes. Previously, ASD subtypes have been defined based on clusters of social communication behaviours and RRBs (Georgiades et al., 2012), structural MRI (Hrdlicka et al., 2005), and various neuroanatomical features (Hong et al., 2017), and FC (Chen et al., 2015). Chen et al. (2015) found two subtypes that differed in terms of ASD symptom severity. Further, Hong et al. (2017) found that prediction of individual scores on the Autism Diagnostic Observation Scale (ADOS) greatly improved when subtypes were considered, compared to considering all ASD participants as one group. Thus, brain-based subtyping has the potential to elucidate brain-behaviour relationships that are unique to each subtype, as it could be the case that certain behaviours result from complex interplay between local and distributed processing in the brain. One limitation of these studies is that they did not include both ASD and TD participants in the subtyping procedures. Given the heterogeneous nature of ASD, the inconsistent reports of FC differences between those with and without ASD, and recent evidence showing a continuum of the relationship between neurobiological features and subclinical ASD symptoms in healthy controls (Rashid et al., 2018), it is crucial to include controls in subtyping analyses as well.

In the present study, we used a data-driven approach to characterize subtypes based on distinct clusters of FC in all participants, and to relate FC patterns to specific behavioural profiles in these subtypes. We first used k-means clustering, an unsupervised machine learning technique, to define subtypes using functional connections as features, and implemented a multivariate statistical analysis that, when applied to neuroimaging data, reveals the optimal relationship between measures of brain activity and experimental design or group membership. This approach allowed us to determine which connections were reliably different between subtypes, and between ASD and TD participants. We also used this multivariate approach to characterize relationships between particular patterns of FC and a set of behaviours. It was hypothesized that defining FC-based subtypes of ASD and TD participants using data-driven metrics would reveal unique information about brain-behaviour interactions.

## RESULTS

### FC-based subtypes of ASD and TD participants

FC-based subtypes were defined using k-means clustering. The effects of scan site were regressed out of the FC data; when these effects were not removed, there was a significant difference in the distribution of scan sites between the two subtypes, *X*^2^(4, *N*=266) = 78.60, *p* <0.001. At this point, subtypes were significantly different in age, *t*(264) = 2.50, *p* = 0.01; thus, effects of both site and age were regressed from the data. As it has been recently shown that despite implementing preprocessing steps that aim to correct for head motion in resting-state fMRI, residual motion effects can contaminate FC estimates (Ciric et al., 2017), a multivariate brain-behaviour analysis was performed to determine if there were relationships between FC and head motion metrics (mean FD and percentage of frames exceeding 0.2mm). There was not a significant relationship between FC and motion (*p* = 0.57).

The optimal number of clusters, as determined by the elbow point criterion, was 2 (Fig. 1A). Using a bootstrapping procedure to evaluate the reliability of the optimal number of clusters, it was found that the optimal number of clusters was 2 in 500/500 bootstrap samples. Subtype 1 consisted of 85 ASD participants and 54 TD participants. Subtype 2 consisted of 60 ASD participants and 67 TD participants. Qualitatively, Subtype 1 was defined by stronger FC between networks, particularly between the DMN and other networks, and weaker FC within networks relative to Subtype 2 (Fig. 1B).

**Fig. 1.**
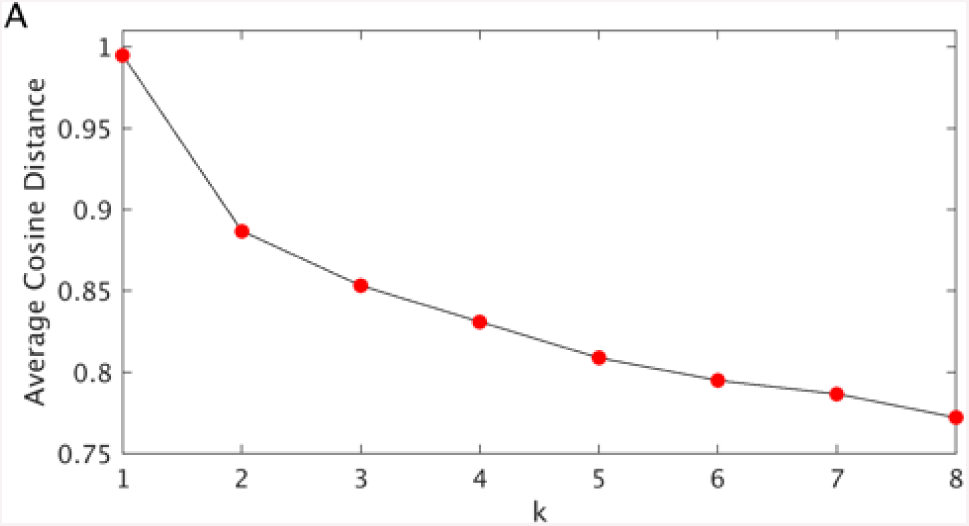
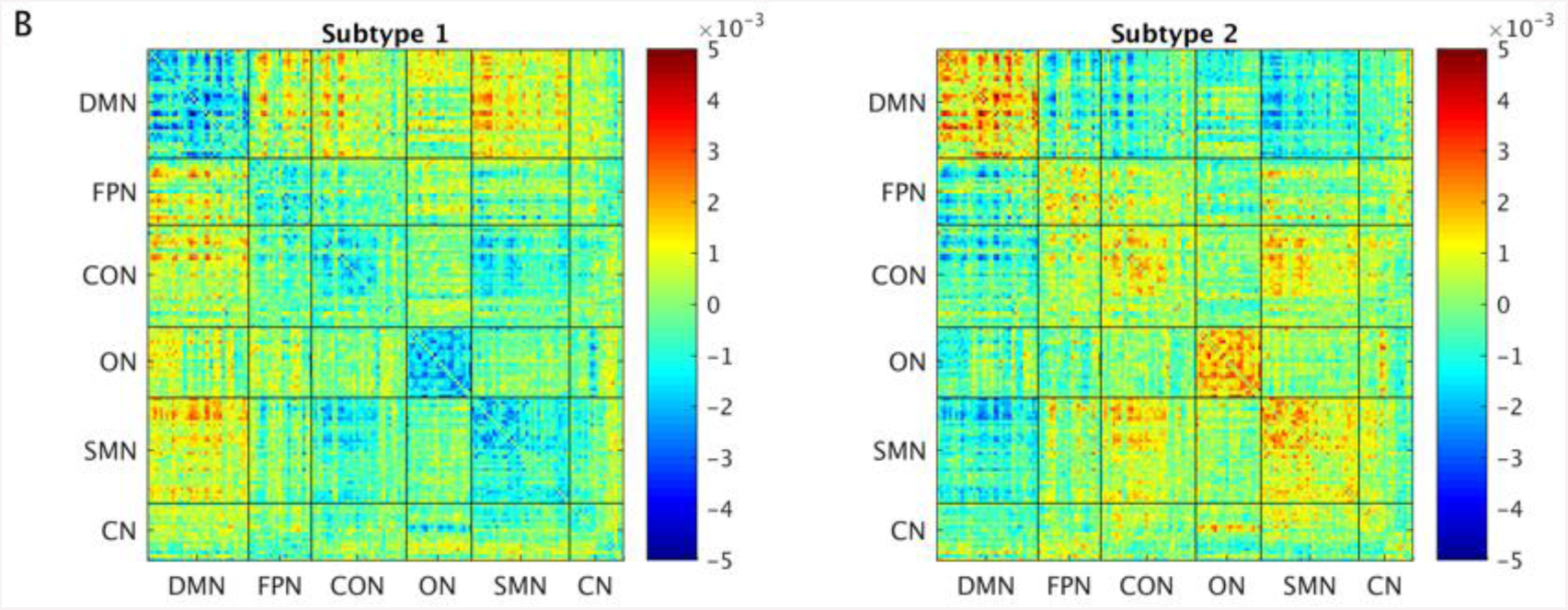
A) Elbow point plots, indicating that the optimal number of clusters is 2. B) Subtype centroids. DMN = default mode network; FPN = frontoparietal network; CON = cingulo-opercular network; ON = occipital network; SMN = sensorimotor network; CN = cerebellar network.

Importantly, subtypes did not differ in demographics or behaviour, including IQ, eye status, medication use, presence of comorbidities, head motion, or the parameters (scan site and age) that were regressed out of the FC matrices (Supplementary Table 3). While subtypes differed in ADOS scores, and differences SRS scores approached significance, these differences were driven by the fact that there were more TD participants with these scores in Subtype 2 compared to Subtype 1. SRS scores did not differ between ASD participants in Subtypes 1 and 2, and also did not differ between TD participants in Subtypes 1 and 2. ADOS scores did not differ between ASD participants in Subtypes 1 and 2, but could not be compared for TD participants in Subtypes 1 and 2, because ADOS scores were only available for 2 TD participants in Subtype 1 and 12 TD participants in Subtype 2.

Next, we used a multivariate statistical approach to determine differences in FC between subtypes and between ASD and TD participants. The reliability of these patterns was determined via bootstrap sampling. A functional connection was considered to be reliable, or stable, if the absolute value of its bootstrap ratio (BSR) exceeded 2. This analysis revealed two significant patterns. The first pattern showed stable differences in FC between subtypes (*p* < 0.001, 61.07% of variance explained, Fig. 2A), whereby Subtype 2 was characterized by stronger FC within resting-state networks, and weaker FC between networks, compared to Subtype 1. The contrast expression for this FC pattern (Supplementary Fig. 2) revealed that functional connections with significant positive BSRs, on average, were positive in Subtype 1 and negative in Subtype 2, and vice versa for negative BSRs. The second pattern revealed a contrast between diagnostic groups (*p* = 0.02, 21.74% of variance explained, Fig. 2B), with a diffuse spatial pattern. The contrast expression for the second pattern (Supplementary Fig. 3) revealed that functional connections with significant positive BSRs, on average, were negative in the ASD group and positive in the TD group, and vice versa for negative BSRs. The third pattern, which revealed a subtype by diagnosis interaction, was not significant, *p* = 0.92. The significance of these spatial patterns within and between resting-state networks (RSNs) was evaluated using permutation tests (see Materials and Methods), and is shown in Fig. 3.

**Fig. 2.**
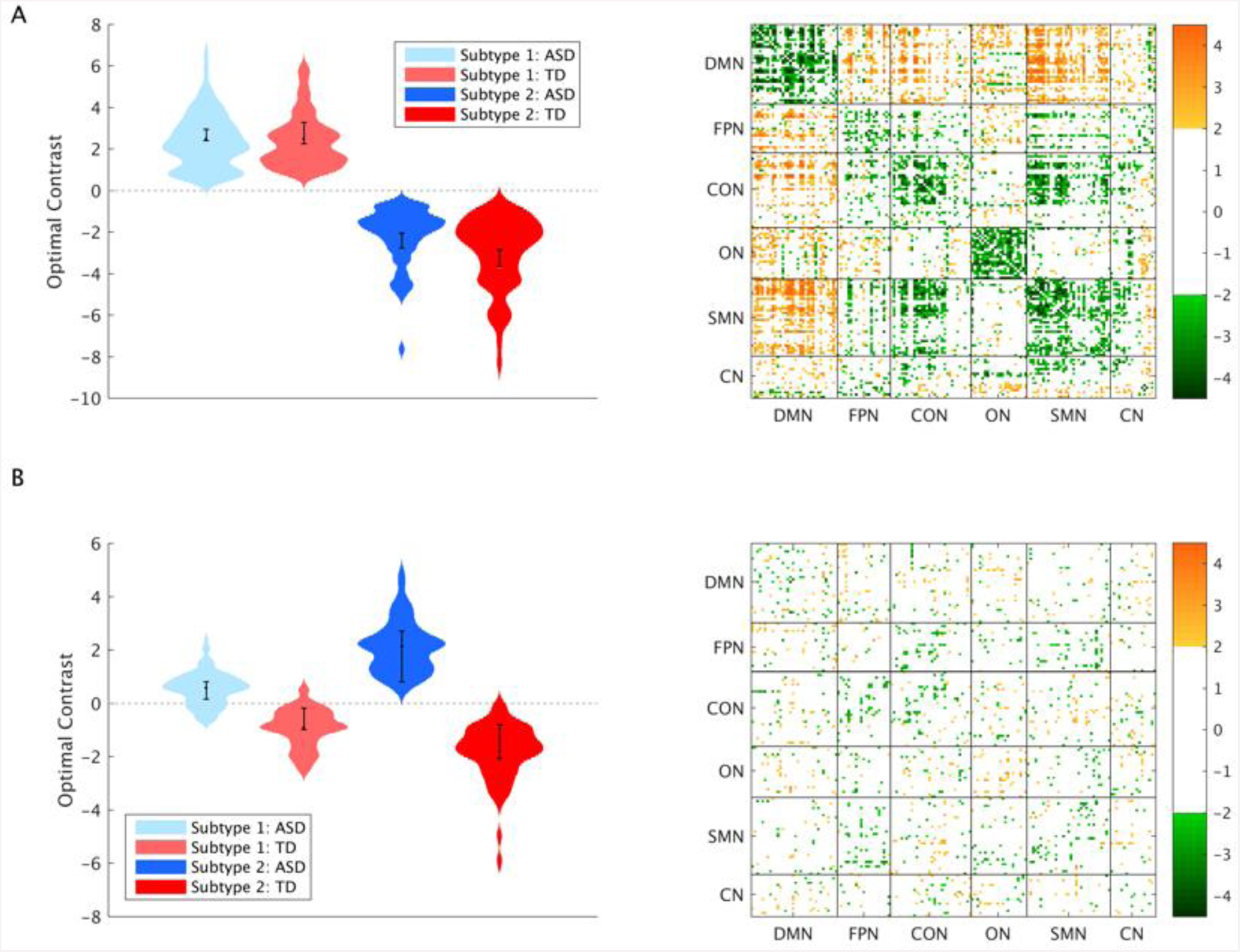
Results from the multivariate group analysis. A) First pattern, and B) second pattern, and the associated BSRs for each connection, at a threshold of ±2. Error bars show 95% confidence intervals determined through bootstrap resampling.

**Fig. 3.**
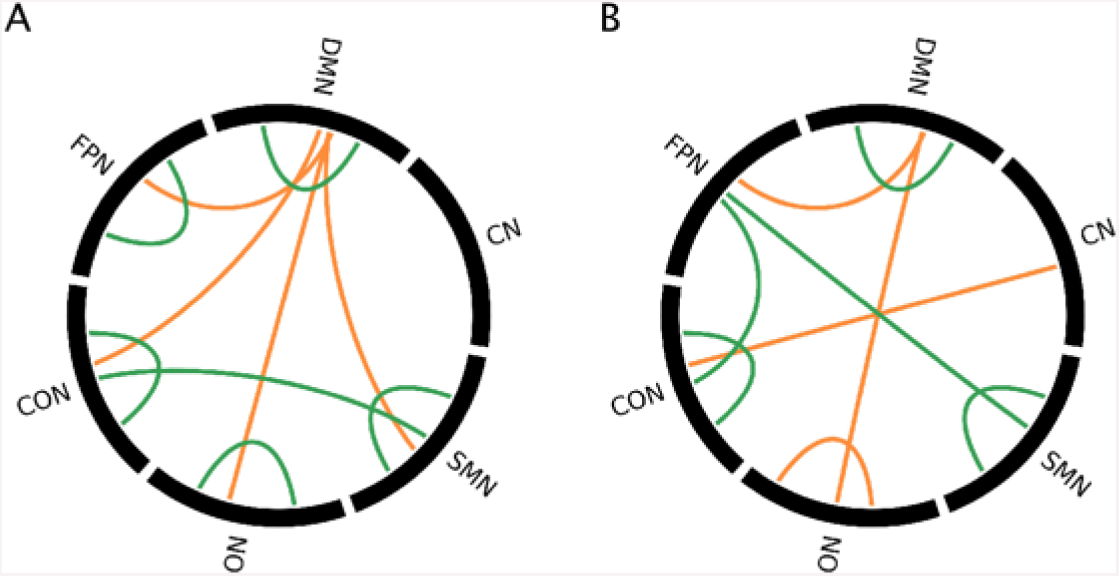
Significant contributions of RSN pairs to each pattern for positive and negative BSRs, for the A) first pattern and B) second pattern from the multivariate group analysis. Orange = positive BSRs, green = negative BSRs.

### Multivariate analyses of FC-behaviour relationships

A multivariate brain-behaviour analysis was used to assess relationships between FC and a set of behavioural measures in the two ASD-TD subtypes, including IQ, ADOS scores (communication (COMM), social affect (SA), and restricted and repetitive behaviours (RRB)), and scores on the Social Responsiveness scale (SRS). The full set of behavioural measures was available for 51 participants (49 ASD, 2 TD) in Subtype 1 and 50 (38 ASD, 12 TD) participants in Subtype 2. ADI-R scores were not included, as only 28 participants in Subtype 1 and 26 participants in Subtype 2 had the full set of behavioural measures including ADI-R scores. Further, none of the participants with the full set of scores including ADI-R scores were TD participants.

The analysis revealed 3 significant patterns. The first pattern (*p* = 0.03, 32.09% covariance explained) revealed stable relationships between FC and IQ and ADOS RRB scores in Subtype 1, and stable relationships between FC and all behavioural measures in Subtype 2. The first brain-behaviour pattern was a contrast between Subtypes 1 and 2 in terms of relationships with FC and ADOS RRB scores, such that connections that were reliably positively correlated with ADOS RRB scores in Subtype 1 were negatively correlated in Subtype 2, and vice versa. The third pattern (*p* = 0.008, 10.82% covariance explained) revealed a different spatial pattern that exhibited stable correlations with IQ and SRS in Subtype 1, and with all ADOS scores and SRS in Subtype 2. Additionally, there was a contrast between Subtypes 1 and in terms of correlations between FC and SRS scores. The seventh pattern (*p* = 0.003, 4.45% covariance explained) revealed a contrast between Subtypes 1 and 2 in terms of correlations between FC and ADOS communication scores, as well as stable correlations between FC and ADOS social affect scores in Subtype 1. Overall, it can be seen that connections that show stable correlations with behaviour are diffuse. Patterns that accounted for more than 10% of the covariance between FC and behaviour (that is, patterns 1 and 3) are shown in Fig. 4, and the corresponding contrast expressions are shown in Supplementary Fig. 4 and 5. The stability of these FC-behaviour relationships within and between RSNs are shown in Fig. 5.

**Fig. 4.**
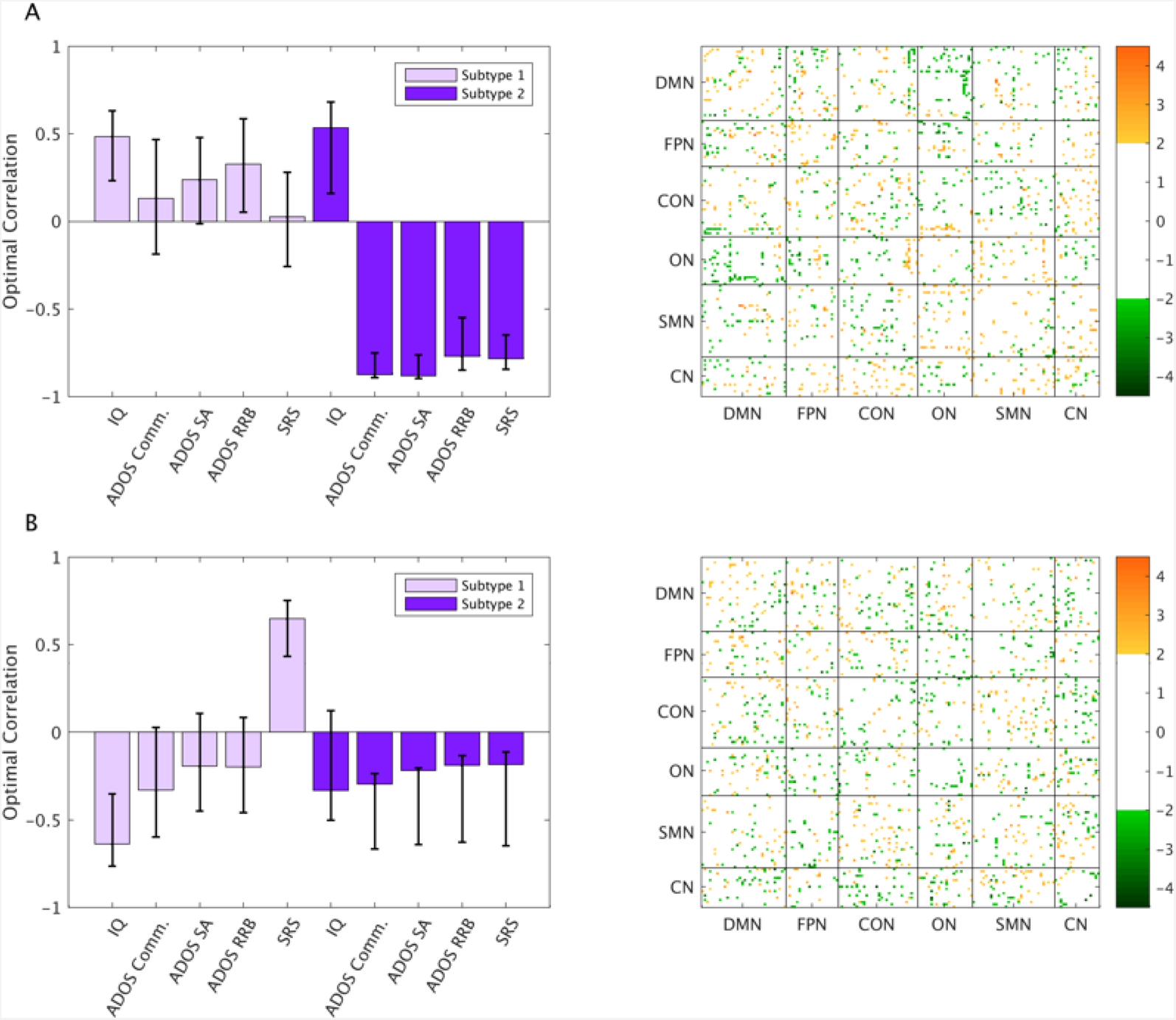
Results from the multivariate brain-behaviour analysis. A) First pattern, and B) third pattern, and the associated BSRs for each connection, at a threshold of ±2. Error bars show 95% confidence intervals determined through bootstrap resampling.

**Fig. 5.**
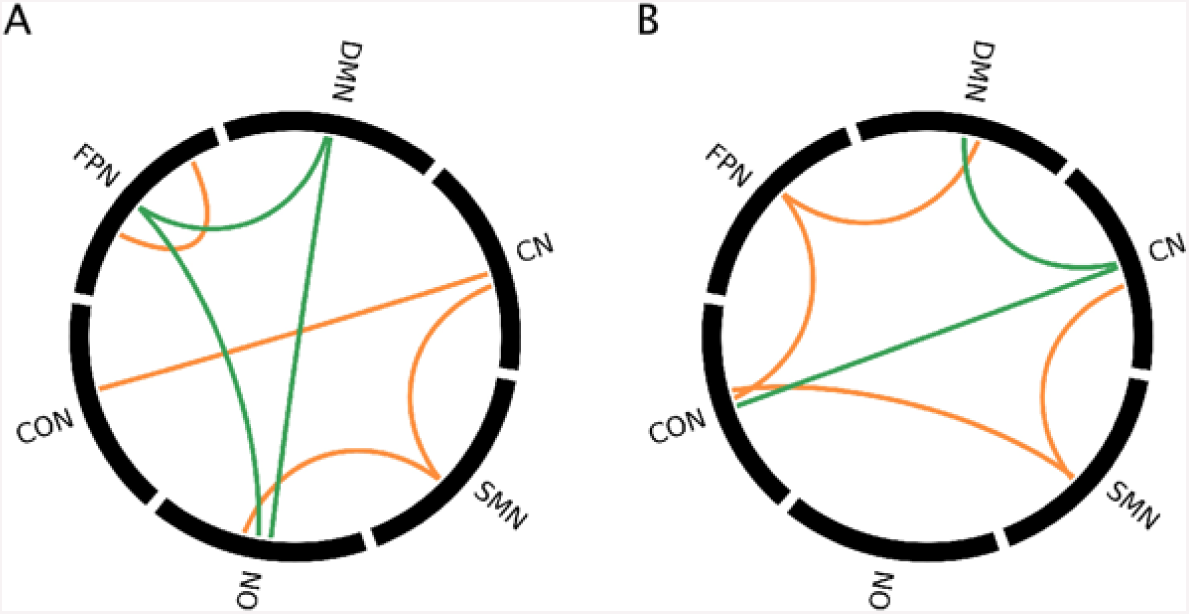
Significant contributions of RSN pairs to each pattern for positive and negative BSRs, for A) first pattern and B) third pattern. Orange = positive BSRs, green = negative BSRs.

The relationship between brain and behaviour scores for ASD and TD participants in Subtype 2 for the first pattern of the multivariate brain-behaviour analysis is shown in Fig. 6. The continuum of scores for both brain and behaviour variables illustrates that there is a pattern of FC that co-varies with the severity of behaviours across the autism spectrum and typical development. This analysis was only performed in Subtype 2, as there were only 2 TD participants in Subtype 1 who had the full set of behaviour measures.

**Fig. 6.**
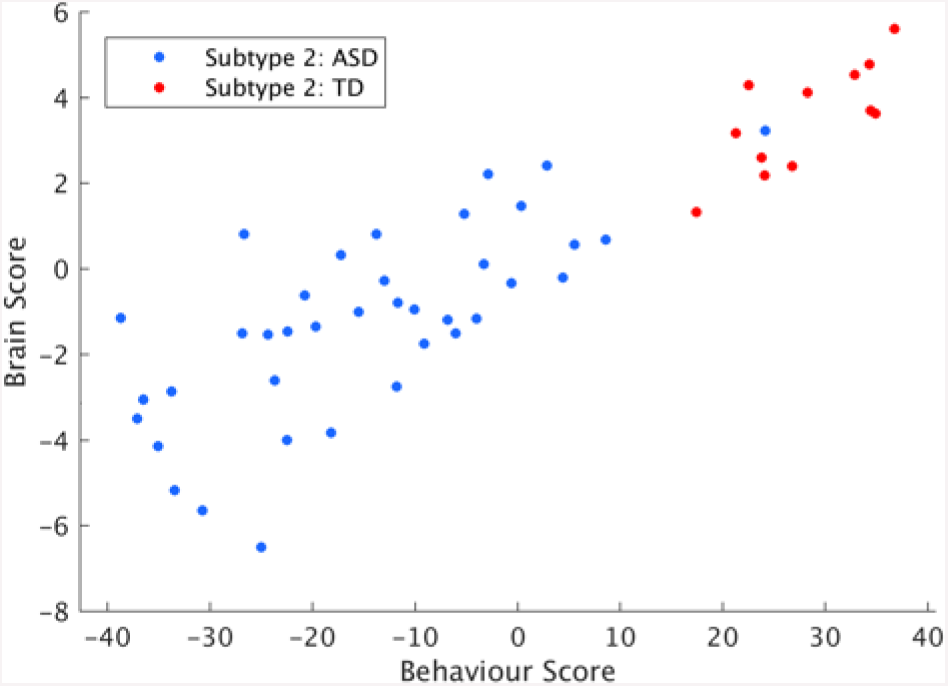
Brain and behaviour scores for Subtype 2, from the first pattern of the multivariate brain-behaviour analysis.

We then determined the relationship between the patterns from the multivariate group analysis and the multivariate brain-behaviour analysis by correlating the brain saliences for each analysis, and evaluated the significance of these correlations using permutation testing. There was a significant correlation between the first brain-behaviour pattern and the second group pattern, *r* = 0.40, *p* < 0.001, indicating that the continuum of FC-behaviour relationships was associated with the diagnostic pattern from the group analysis. The correlations between the other patterns were not significant: (brain-behaviour pattern 1 and group pattern 1: *r* = −0.06, *p* = 0.81; brain-behaviour pattern 3 and group pattern 1: *r* = 0.005, *p* = 0.45; brain-behaviour pattern and group pattern 2: *r* = 0.07, *p* = 0.13).

## DISCUSSION

### Overview

This study reveals that there are distinct clusters of FC patterns in both ASD and controls. We characterized network-level differences between subtypes and diagnostic groups, and further showed that individuals within each subtype exhibit different relationships between FC metrics and behavioural measures. The continuum of brain and behaviour scores across ASD and TD participants reveals that FC phenotypes observed in ASD extend to typical development in relation to behavioural severity.

### Comparison of FC between subtypes and diagnostic groups

Two FC-based subtypes were defined for all participants. When all four groups were considered in a multivariate analysis (i.e. ASD Subtype 1, ASD Subtype 2, TD Subtype 1, and TD Subtype 2), the strongest pattern, not surprisingly, was a contrast between subtypes. Regardless of diagnostic group, Subtype 2 was defined by greater FC within networks and lower FC between networks, especially between the DMN and other RSNs, compared to Subtype 1. Connections within networks tended to be positive on average in Subtype 2 and negative in Subtype 1, indicating reduced interactions among brain regions within these networks in Subtype 1. Further, connections between networks that were lower in Subtype 2 tended to be negative, but were positive on average in Subtype 1 (Supplementary Fig. 2). As anti-correlations between resting-state networks are hypothesized to signify the division of labour between brain regions that are involved in different functions (Fransson 2006), and the ability for regions that are relevant for certain cognitive functions to become activated with concurrent deactivation of irrelevant regions (Fox et al., 2005; Greicius et al., 2003), these abilities may be affected in Subtype 1.

A second pattern revealed diffuse functional connections that differed between diagnostic groups in both subtypes. ASD participants exhibited reliable decreases in FC within the SMN, DMN and CON, but greater FC within the ON. Atypical FC of sensorimotor regions in ASD has reported in previous studies (Anderson et al., 2011; Mostofsky et al., 2009; Turner et al., 2006). Thus, despite the broad range of sensorimotor difficulties in ASD (Minshew et al., 1997; Perry et al., 2007; Whyatt & Craig, 2013), atypical SMN FC may be common across the autism spectrum. It has been hypothesized that abnormal DMN functioning in ASD relates to decreased self-referential processing, decreased abilities to redirect attention from external to internal processing, and difficulties with theory of mind (e.g. Assaf et al., 2010). Various studies have reported decreased FC between DMN regions in ASD (Assaf et al., 2010; Kennedy & Courchesne, 2008; Monk et al., 2009; Weng et al., 2010), although hyperconnectivity has also been reported (Monk et al., 2009; Uddin et al., 2013a). Decreased FC within the CON, which plays a role in stable set-maintenance (Dosenbach et al., 2007), is line with previous studies that showed difficulties with set-maintenance in ASD (Kaland, Smith, & Mortensen, 2008; Miller et al., 2015). Increased FC in the ON is consistent with findings of increased local connectivity in primary visual regions (Keown et al., 2013) and increased involvement of extrastriate cortex (Shen et al., 2012) in ASD. Elevated FC in right ventral occipital-temporal cortex in ASD has been associated with higher social deficits (Chien et al., 2015). Additionally, reliably higher FC was found between the DMN and FPN, DMN and ON, and CON and CN in ASD participants. These connections were positive on average in ASD, but negative on average in controls (Supplementary Fig. 3). Previous studies have also reported reduced negative connectivity in ASD, which was described as reduced functional segregation of networks (Rudie et al., 2012; 2013a). However, other between-network connections (FPN-CON and FPN-SMN) exhibited a greater degree of anti-correlation in ASD. The functional significance of decreased anti-correlations between some resting-state networks, but increased anti-correlations between others, remains to be explored.

The third pattern, showing a subtype by diagnosis interaction, was not significant, thus revealing additive effects of subtype and diagnosis on FC patterns. Thus, the expression of the subtypes does not depend on the diagnosis; the manifestation of the subtypes in ASD is not different from controls.

### Comparison of FC-behaviour relationships between subtypes

Reliable correlations between FC and behaviour were observed both within and between RSNs for IQ and ADOS RRB scores for Subtype 1, and all behavioural measures for Subtype 2, showing that similar behavioural profiles can be associated with different functional correlates in the brain. Previous studies have reported mixed results regarding associations between FC measures and ASD behavioural measures. For instance, Keown et al. (2013) found that overconnectivity in posterior brain regions was associated with greater severity ASD severity, and that frontal underconnectivity was found only in low-severity participants. However, another study found that ASD severity was correlated with the extent of hyperconnectivity in the salience network, which includes regions such as the dorsal anterior cingulate cortex and frontoinsular cortex (Uddin et al., 2013b). Lee et al. (2016) reported overall reduced FC density in ASD, and found that average interhemispheric FC density and contralateral FC density in a lingual/parahippocampal gyrus cluster and default mode network regions was negatively correlated with RRBs. On the other hand, hyperconnectivity between the posterior cingulate cortex (PCC), a core region of the DMN, and the right parahippocampal gyrus was associated with more severe RRBs in another study (Monk et al., 2009). Our results highlight the importance of considering FC-based subtypes when examining brain-behaviour relationships in individuals with and without ASD. Importantly, individuals in each subtype did not differ significantly in IQ or SRS scores, and ASD participants in the two subtypes did not differ significantly in ADOS scores. Thus, there is unique information about FC-based subtypes that is not accessible by using behaviour alone.

The multivariate brain-behaviour analysis supports the idea that instead of being a categorical diagnosis, ASD should indeed be considered as an extreme of a continuum of both neurobiological and behavioural features that can also be observed in TD individuals (Constantino & Todd, 2003; Rashid et al., 2018). In other words, there is normal variation in FC across both ASD and TD participants (see Fig. 6), but too much of this natural variation is associated with a diagnosis of ASD. This idea is supported by the continuum of brain and behaviour scores from pattern 1 of the brain-behaviour analysis for Subtype 2, and the significant correlation between the spatial pattern for this pattern and the second pattern from the group analysis, that is, the contrast in FC between diagnostic groups. This dimensional approach has also been reinforced by recent studies that reported novel findings in individuals with ASD by accounting for the heterogeneity of the relationships between behavioural severity and various neurobiological features (Hahamy et al., 2015; Nunes et al., 2018). Recently, it has been pointed out that different features of brain function are variable even among TD individuals, and a certain feature cannot be considered to be an impairment unless it is accompanied by behavioural symptoms (Muller & Amaral, 2017). Our results support this idea by showing that some sets of functional connections are a) similar among subsets of ASD and TD participants, and b) correlated with behavioural severity. The similarity of FC patterns in ASD and controls has also been demonstrated in a recent study by Spronk et al. (2018), which demonstrated that resting-state FC patterns between TD participants and several clinical groups, including ASD, attention deficit hyperactivity disorder, and schizophrenia, are highly correlated, despite the presence of clinical symptoms.

### Limitations

One limitation of our study is that we defined subtypes using a single data preprocessing strategy. It has been proposed that differences in analysis approaches between studies are the most likely causes of inconsistent results between studies of FC in ASD (Hull et al., 2016). For instance, it has been shown that global signal regression reduces the relationship between FC and head motion, but can result in distance-dependent artifacts in FC unless used in combination with censoring methods (Ciric et al., 2017). Preprocessing strategies such as global signal regression and low-pass filtering have been shown to affect group differences in FC between participants with and without ASD (Gotts et al. 2013; Muller et al., 2011). The length of fMRI scans may also contribute to heterogeneity across studies: it has been suggested that increasing scan lengths, for instance from 5 to 13 minutes, improves the reliability of FC estimates (Birn et al., 2013). It is therefore crucial to gain a better understanding of how preprocessing choices and scanning parameters affect group differences in FC, and to compare FC-based subtypes across different preprocessing strategies.

Subtypes in this study were defined based on FC. The incorporation of additional metrics may help to further characterize differences between the two subtypes defined in this study. For instance, recent work has focused on altered dynamic FC “states” in ASD (e.g. Chen et al., 2017; de Lacy et al., 2017; Rashid et al., 2018). However, as participants’ time series consisted of only 145 time points, characterizing FC states in this dataset was not feasible.

Finally, we examined the continuum of brain and behaviour scores across both ASD and TD participants in Subtype 2; however, ADOS scores were available for only 2 TD participants in Subtype 1. It will be important for future studies to collect ADOS scores in TD participants to better characterize the continuum of FC-behaviour relationships across all participants in multiple subtypes.

### Conclusions

Multivariate analyses of FC-based subtypes highlight the importance of considering the heterogeneity of FC patterns and measures of behaviour in resting-state studies, and reveal the continuum of brain-behaviour relationships in individuals with and without ASD. As subtypes exhibited different relationships between FC and behavioural severity, it will be important to determine if individuals with ASD in different subtypes exhibit unique responses to treatments and behavioural therapies.

## MATERIALS AND METHODS

### Participants

Resting-state fMRI data from 145 males with ASD and 121 TD males were acquired from the Preprocessed Connectomes Project (PCP; Craddock et al., 2015; http://www.preprocessed-connectomes-project.org/abide). The data had been obtained from the Autism Brain Imaging Data Exchange (ABIDE; Di Martino et al., 2014; http://www.fcon_1000.projects.nitrc.org/indi/abide) and preprocessed using the Connectome Computation System pipeline (Xu et al., 2015). Participants were excluded if their age was greater than 40, full scale IQ was less than 75, mean framewise displacement (FD) during the resting-state fMRI scan was greater than 0.20mm, percentage of data points exceeding 0.20mm was greater than 20%, and/or scans were rated as good by less than 2 (out of 3 raters) as per the ABIDE quality assessment protocol (http://preprocessed-connectomes-project.org/abide/quality_assessment.html). Groups were matched for age, IQ, mean framewise displacement and the percentage of data points exceeding 0.20mm. ASD diagnoses were confirmed using the Autism Diagnostic Observation Scale (ADOS; Lord et al., 2000) and/or the Autism Diagnostic Interview-Revised (ADI-R; Lord et al., 1994). Participant characteristics are shown in Table 1, along with the number of scores that were available for ADOS, ADI-R and SRS scores for ASD participants if these scores were not available for all 145 participants. Participant characteristics for each site are described in Supplementary Table 1.

**Table 1:**
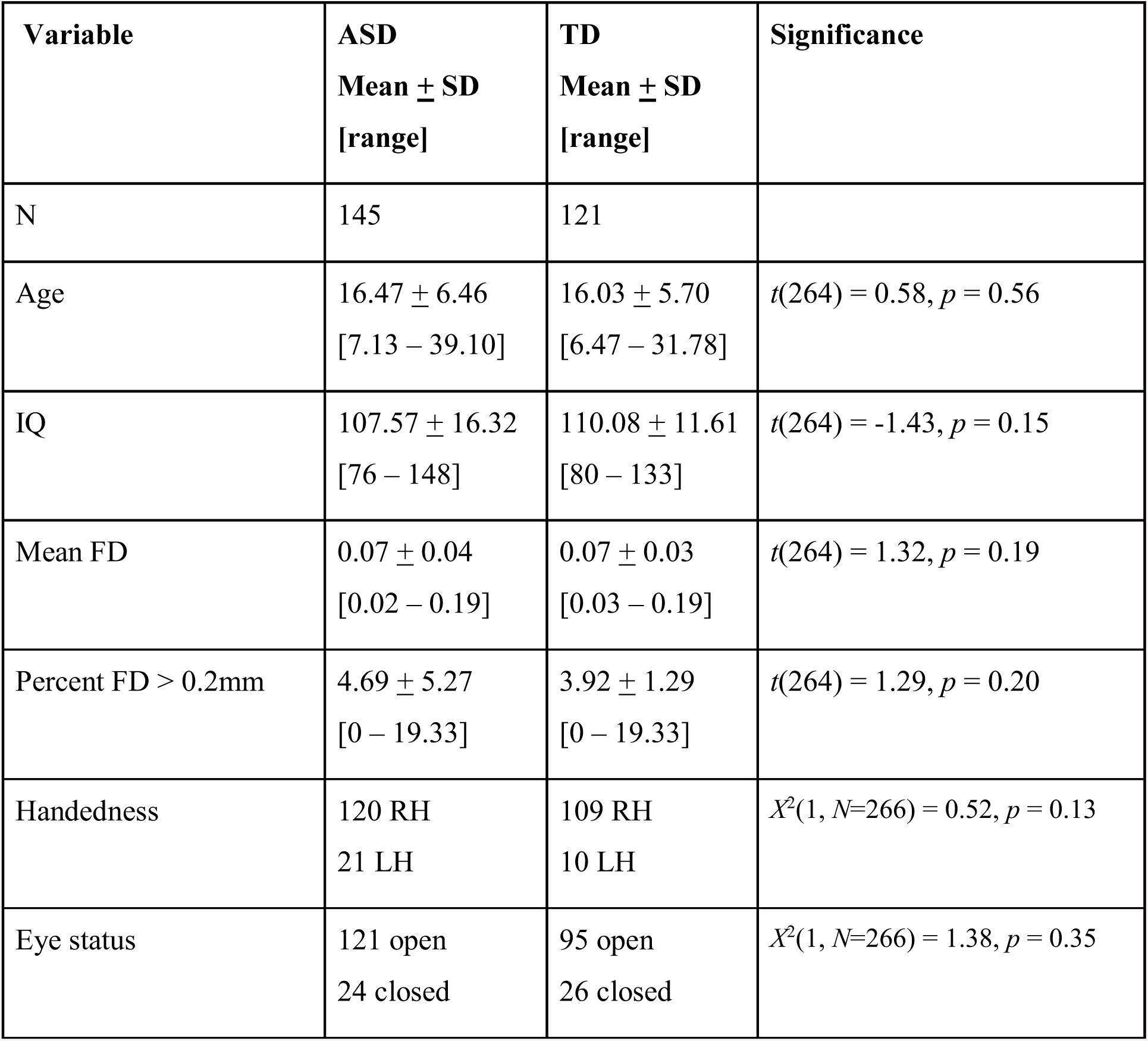
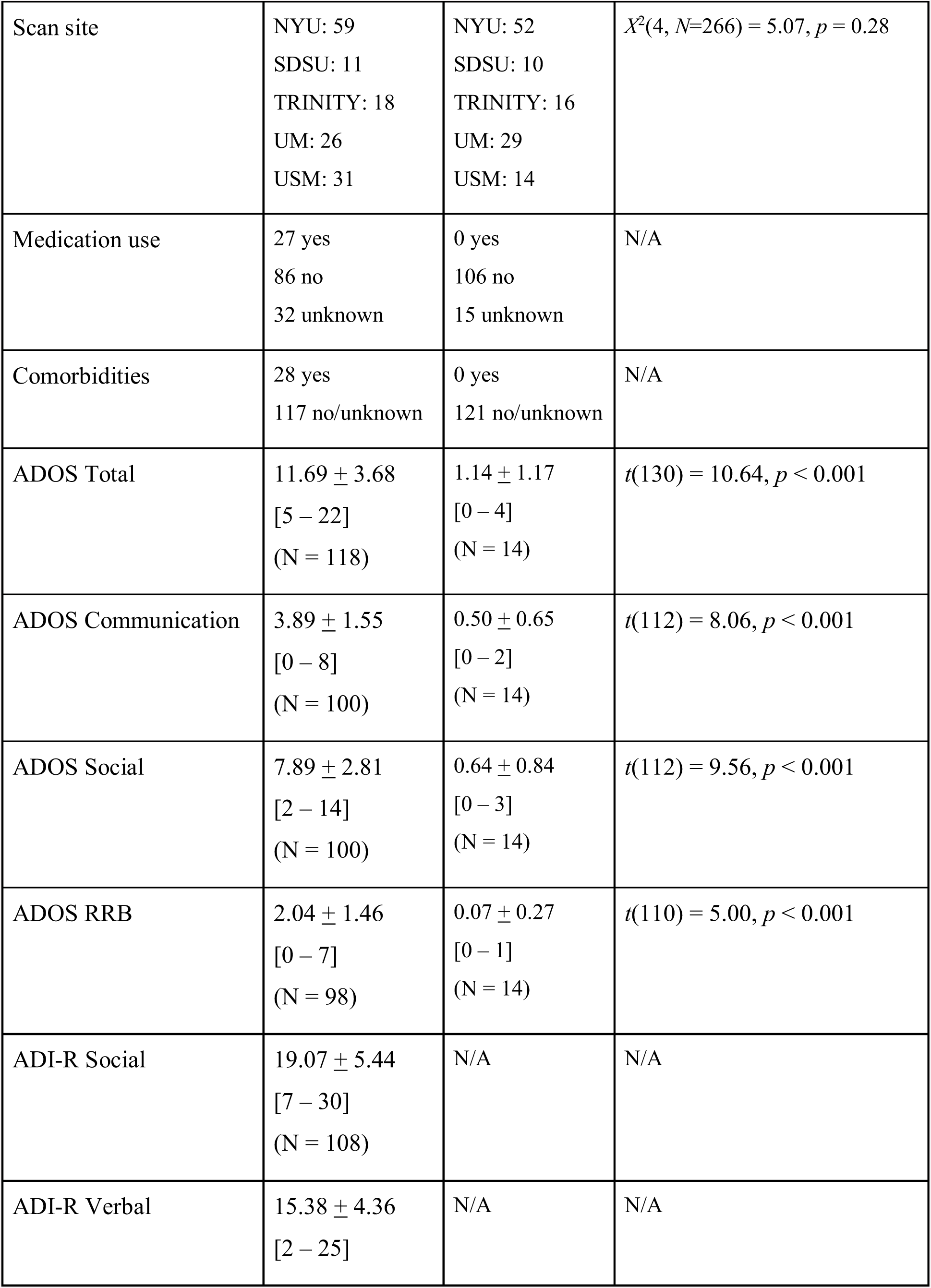
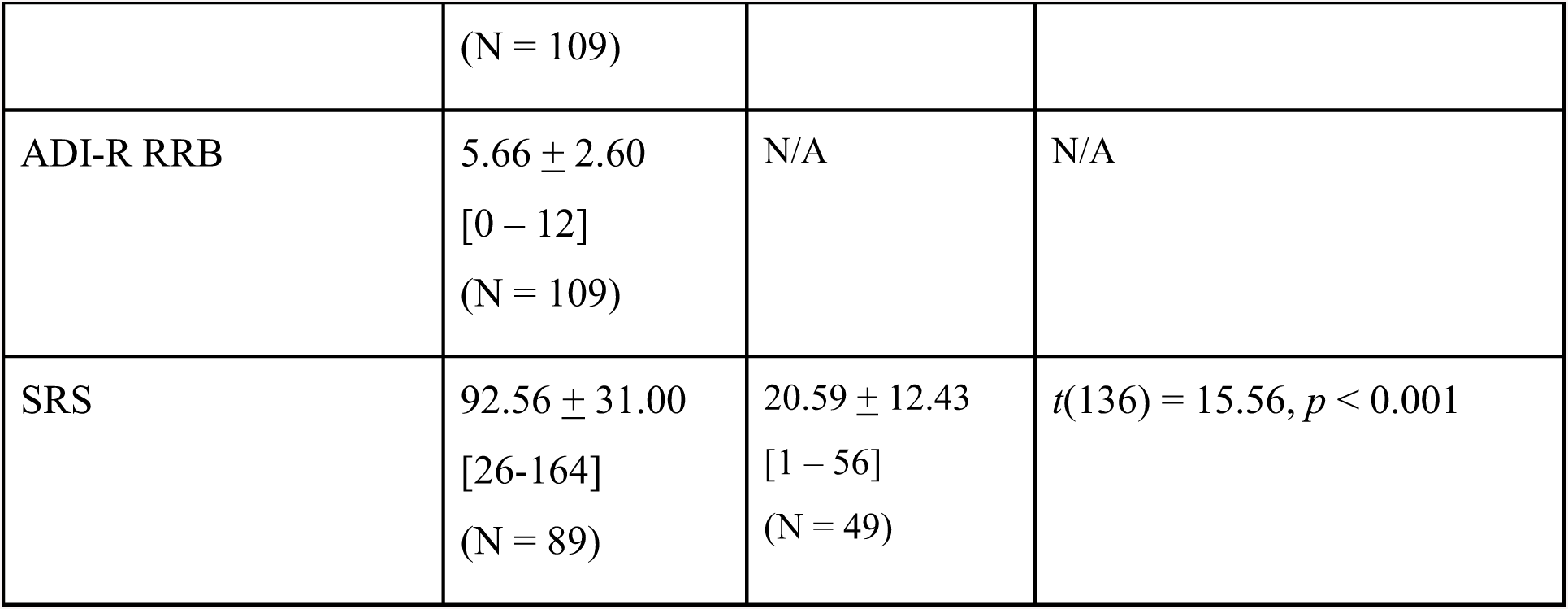
Participant characteristics

### fMRI Preprocessing

Data from five sites (New York University Lagone Medical Center, University of Utah School of Medicine, San Diego State University, Trinity Centre for Health Sciences, and University of Michigan) using a TR of 2000ms were included. The proportion of ASD compared to TD subjects was not significantly different across sites, *X*^2^(4, *N*=266) = 5.07, *p* = 0.28. Written informed consent or assent was obtained for all participants in accordance with respective institutional review boards. Additional information about scanner types and parameters can be found on the ABIDE website (http://www.fcon_1000.projects.nitrc.org/indi/abide). The CCS preprocessing steps, which had been carried out as part of the Preprocessed Connectomes Project, were as follows: dropping the first 4 volumes, removing and interpolating temporal spikes, slice timing correction, motion correction, brain mask creation, 4D global mean-based intensity normalization, boundary-based registration of functional to anatomical images, anatomical segmentation of grey matter, white matter and cerebrospinal fluid, nuisance parameter regression (including 24 motion parameters, white matter and CSF signals, linear and quadratic trends, and the global signal), bandpass filtering (0.01 to 0.1Hz), and registering functional images to the MNI template. The final preprocessed time series for each subject were obtained from the Preprocessed Connectomes Project. We chose to use data that had the global signal regressed out, as this step has been shown to help mitigate differences across multiple sites (Power et al., 2014). Further, it has been shown recently that global signal regression attenuates artefactual changes in BOLD signal that are introduced by framewise displacement (Byrge & Kennedy, 2017). It should also be noted that without global signal regression, FC-based subtypes differed in head motion (both mean framewise displacement (*p* < 0.001) and percentage of frames above 0.2mm (*p* < 0.001). The time series of 160 4.5mm spherical regions of interest (ROIs) from the Dosenbach atlas (Dosenbach et al., 2010) were obtained (see Supplementary Table 2 and Supplementary Fig. 1). Regions in this atlas were selected from meta-analyses of task-related fMRI studies and categorized into six different resting-state networks (RSNs): the default mode network (DMN), frontoparietal network (FPN), cingulo-opercular network (CON), occipital network (ON), sensorimotor network (SMN), and cerebellar network (CN). Additional details of the fMRI preprocessing steps can be found on the PCP website (http://www.preprocessed-connectomes-project.org/abide).

### Functional connectivity

Each subject’s fMRI time series was truncated to 145 time points, which was the minimum number of time points across subjects. FC was defined by Fisher z-transformed Pearson correlations for each ROI pair across all time points for each participant. The effects of age and acquisition site (represented as a Helmert basis) were regressed out of the FC matrices.

### K-means Clustering

K-means clustering was used to define subtypes distinct FC patterns. The lower triangle of each participant’s FC matrix was used, such that the matrix for k-means was in the form subjects x FC. The k-means algorithm begins with an initialization of *k* centroids. Then, in the *assignment* step, each participant is assigned to the closest centroid using the cosine distance, defined as one minus the cosine of the included angle between each subjects’ FC values and each cluster’s centroids, which are treated as vectors. Next, in the *centroid update* step, new centroids are defined as the mean of the data points that are currently assigned to that centroid. These two steps are repeated iteratively until convergence, when cluster assignments no longer change.

The “elbow point” criterion was used to determine the optimal number of clusters. To determine the elbow point, the average cosine distance between a cluster’s centroids and the FC values of participants assigned to that particular cluster is calculated for each cluster, then averaged across clusters to obtain a single distance metric for each value of k. These distances are then plotted as a function of k, and the “elbow” is defined as the value of k where the change in the rate of decrease in distance is sharpest. Values from k = 2 to k = 8 were tested (but also included k = 1 in the elbow point plot as a reference point). Further, we evaluated the reliability of the number of clusters using bootstrap resampling. Fifty percent of the sample was selected at random, and were grouped into subtypes using the k-means algorithm for values of k from 2 to 8. The elbow criterion was then used to select the ideal value of k for the bootstrap sample. This process was repeated 500 times to determine the reliability of the optimal number of clusters.

### Partial Least Squares

Partial least squares (PLS) is a multivariate statistical technique that is used to optimally relate brain activity to experimental design or group membership in the form of latent variables (McIntosh et al., 1996; McIntosh & Lobaugh, 2004; Krishnan et al., 2011). PLS software, which is implemented in Matlab, is available for download from research.baycrest.org/pls-software. In *mean-centering PLS*, patterns relating a matrix of brain variables (in the form subjects x brain variables) and group membership are calculated. For this study, the brain variables were the FC values in the lower triangle of each subject’s FC matrix (12720 connections). Mean-centering PLS was used to examine differences in FC between subtypes and between ASD and TD participants.

Using singular value decomposition (SVD), orthogonal patterns that express the maximal covariance between the brain variables and group membership are computed. The resulting patterns are sorted in order of the proportion of covariance between the brain and design/behaviour variables that the pattern accounts for, with the first pattern accounting for the most covariance. Each pattern consists of saliences (weights) and a singular value. The brain saliences indicate which brain variables (in this case, functional connections) best characterize the relationship between the brain variables and group differences. Design saliences indicate the group differences profiles that best characterize this relationship. Singular values indicate the proportion of covariance between the brain and design matrices that each pattern accounts for. Brain scores, which represent each subject’s contribution to each brain salience, are calculated by multiplying the original matrix of brain variables by the brain salience.

In *behaviour PLS*, a matrix of behaviour variables is also included in the analysis to determine design-dependent (in this case, group-dependent) relationships between the brain variables and behaviour. For this study, behavioural PLS was used to examine associations between FC and a set of behavioural variables including IQ, ADOS scores (communication, social affect, and RRBs), and scores on the Social Responsiveness Scale (SRS) in each subtype.

The statistical significance of each pattern was determined using permutation testing. For this procedure, the rows (participants) of the matrix of brain variables are reshuffled, and new singular values are obtained using SVD. In this study, this procedure was repeated 1000 times to create a distribution of singular values. The p-value associated with the original singular value is defined as the proportion of singular values from the sampling distribution that are greater than the original singular value, thus representing the probability of obtaining a singular value larger than the original value under the null hypothesis that there is no association between the brain variables and group membership.

In addition to determining the statistical significance of each pattern, the reliability of the brain saliences can also be determined by utilizing a bootstrapping procedure. Bootstrap samples are generated by randomly sampling subjects with replacement, while ensuring that group membership is maintained. In this study, 500 bootstrap samples were generated. Creating bootstrap samples allows one to determine which brain variables are stable, regardless of which participants are included in the analysis. The bootstrap ratio (BSR), defined as the ratio of the brain salience to the standard error of the salience (as estimated by the bootstrap procedure), is a measure of this stability. Reliable connections were defined as those that surpassed a BSR threshold of ±2.0, which corresponds roughly to a 95% confidence interval.

As FC values can take on positive or negative values, positive BSRs could correspond to either stronger positive or weaker negative connectivity in one group compared to the other, and negative BSRs could indicate weaker positive or stronger negative connectivity. Thus, expressions of FC PLS contrasts were generated for each group. Positive expressions were generated by averaging connections (Fisher z-transformed Pearson correlation coefficients) that had BSRs greater than 2 across all participants in each group. A similar procedure was performed for negative expressions, that is, for connections showing BSRs less than −2.

In addition to assessing the contribution of each individual connection to the group differences, we were interested in determining the extent to which network-level FC, both within and between RSNs, contributed to the group differences. This was of particular interest due to hypotheses that ASD may be characterized by atypical FC within and between networks (e.g.Hull et al., 2016; Rudie & Dapretto, 2013b). To assess the relative contributions of each RSN to the spatial patterns, the BSR-thresholded spatial maps (i.e. adjacency matrices in the form connections x connections) were separated into positive BSRs and negative BSRs. These maps were thresholded such that connections with a BSR less than 2 but greater than −2 were set to 0. Positive BSRs greater than 2 were set to 1, and negative BSRs less than −2 were set to −1. All thresholded BSRs within each pair of networks were then averaged to obtain a 6×6 matrix showing the average contribution of each network pair to the spatial pattern, separately for positive and negative BSRs. To assess the significance of these contributions, the order of connections in the BSR thresholded matrices was permuted while keeping the RSN labels the same, and then the above procedure was repeated to calculate the RSN contributions. This process was repeated 1000 times to obtain a distribution of average contribution values for each RSN pair. Then, the significance of the original contribution is defined as the proportion of contribution values from the sampling distribution that are greater than or equal to the original value.

#### 2.6. Data visualization

Connectivity circle plots were created using the plot_connectivity_circle function from the open-source MNE software package implemented in Python (Gramfort et al., 2013; 2014). All other figures were created using Matlab (MATLAB 8.6.0 (R2015b), MathWorks, Natick, MA). Violin plots were created using the distributionPlot.m function (Jonas 2017).

## ACKNOWLEDGEMENTS

The authors thank Bratislav Misic and Sam Doesburg for helpful discussions, and the contributors to the Autism Brain Imaging Exchange and Preprocessed Connectomes project.

This work was supported by an Ontario Graduate Scholarship (OGS), Mynne & Harold Soupcoff Fellowship, and Finkler Graduate Student Fellowship to A.K. Easson.

## Abbreviations

ABIDE: Autism Brain Image Data Exchange
ADI-R: Autism Diagnostic Interview Revised
ADOS: Autism Diagnostic Observation Scale
ASD: autism spectrum disorder
BSR: bootstrap ratio
CN: cerebellar network
COMM: communication
CON: cingulo-opercular network
Cov.= covariance; DMN: default mode network
FC: functional connectivity
FD: framewise displacement
FPN: frontoparietal network
ON: occipital network
PCP: Preprocessed Connectomes Project
PLS: partial least squares
ROI: region of interest
RRBs: restricted and repetitive behaviours
RSN: resting-state network
SA: social affect
SMN: sensorimotor network
SRS: Social Responsiveness Scale
SVD: singular value decomposition
TD: typically developing

